# Polymorphism-aware estimation of species trees and evolutionary forces from genomic sequences with RevBayes

**DOI:** 10.1101/2021.12.15.472751

**Authors:** Rui Borges, Bastien Boussau, Sebastian Höhna, Ricardo J. Pereira, Carolin Kosiol

## Abstract

1. The availability of population genomic data through new sequencing technologies gives unprecedented opportunities for estimating important evolutionary forces such as genetic drift, selection, and mutation biases across organisms. Yet, analytical methods that can handle polymorphisms jointly with sequence divergence across species are rare and not easily accessible to empiricists.
2. We implemented polymorphism-aware phylogenetic models (PoMos), an alternative approach for species tree estimation, in the Bayesian phylogenetic software RevBayes. PoMos naturally account for incomplete lineage sorting, which is known to cause difficulties for phylogenetic inference in species radiations, and scale well with genome-wide data. Simultaneously, PoMos can estimate mutation and selection biases.
3. We have applied our methods to resolve the complex phylogenetic relationships of a young radiation of *Chorthippus* grasshoppers, based on coding sequences. In addition to establishing a well-supported species tree, we found a mutation bias favoring AT alleles and selection bias promoting the fixation of GC alleles, the latter consistent with GC-biased gene conversion. The selection bias is two orders of magnitude lower than genetic drift, validating the critical role of nearly neutral evolutionary processes in species radiation.
4. PoMos offer a wide range of models to reconstruct phylogenies and can be easily combined with existing models in RevBayes — e.g., relaxed clock and divergence time estimation — offering new insights into the evolutionary processes underlying molecular evolution and, ultimately, species diversification.

## Introduction

The recent development of sequencing technologies has made it possible to obtain large amounts of genomic data at a low cost, across model and non-model organisms. We now have an unprecedented opportunity to study macroevolutionary and microevolutionary processes at temporal scales where the distinction between species and population is challenging to establish (Leaché and Oaks, 2017). To resolve the intricate phylogenetic relationships between closely related taxa, we need models of species divergence that are able to assess the significance of different evolutionary forces in shaping species’ diversity patterns. The polymorphism-aware phylogenetic models (PoMos) are examples of such models (De Maio et al., 2013, 2015). PoMos describe the evolutionary relationship between taxa (i.e., populations or species), in which the frequency of alleles change over time, conditioned on evolutionary forces such as mutational bias, genetic drift, and selection (Borges et al., 2019). Thus, PoMos have the ability to leverage population genomic data to estimate parameters of molecular evolution and jointly provide more accurate estimates of the species tree.

PoMos can be seen as an extension of the traditional phylogenetic substitution models (e.g., JC, HKY, and GTR) (Jukes and Cantor, 1969; Hasegawa et al., 1985; Tavaré, 1986), in the sense that PoMos additionally consider the existence of polymorphisms. Traditional phylogenetic substitution models assume that a species can be represented using a single genotype and that all differences occur only between (but not within) species. By describing allele changes over time, PoMos operate in a state-space that includes both fixed and polymorphic states. The possibility to model the existence of ancestral polymorphisms is of particular importance as PoMos naturally accounts for incomplete lineage sorting — i.e., the persistence of ancestral polymorphisms during speciation —, which is known to be a prime cause of phylogenomic conflict in species tree inference (Maddison and Knowles, 2006; Szöllősi et al., 2015).

PoMos are similar to the multispecies coalescent approach because both allow estimating the species tree. However, PoMos describe allele trajectories instead of genealogies, therefore integrating over all possible genealogical histories to directly estimate the species tree (similar to SNAPP, Bryant et al., 2012) and bypassing the computational burden of estimating genealogies. This burden continues to be a major constraint for the coalescent-based species tree methods, as the space of unknown genealogies is large and difficult to explore (Ogilvie et al., 2016; Yang and Rannala, 2017; Flouri et al., 2020). The integration over genealogies in PoMos allows them to scale well with genomic data sampled from many populations and individuals (Schrempf et al., 2016). The neutral PoMos were previously integrated into the maximum likelihood framework in IQ-Tree, primarily for species tree inference (Schrempf et al., 2019). PoMos were previously integrated into the maximum likelihood and Bayesian framework respectively in IQ-Tree (Schrempf et al., 2019) and BEAST 2 Bouckaert et al. (2019), primarily for species tree inference. However, an implementation including new PoMo models, like the recently proposed virtual PoMos and models with selection and different mutation schemes is still missing Borges et al. (2019, 2022). More importantly, an easy-to-expand and well-documented implementation of PoMos is needed. The implentation of PoMos in RevBayes opens possibilities of combining PoMos with other sophisticated phylogenetic models including molecular dating, phylogeography, trait evolution, and branch-heterogeneous processes, and will permit tackling evolutionary questions that are previously intractable.

Furthermore, PoMos permit disentangling the contribution of the various evolutionary forces to the species divergence process that are often confounded in existing methods. For example, selection favoring the fixation of specific alleles can be particularly difficult to model under the coalescent approaches but it is quite straightforward to do under PoMos (Borges et al., 2019). PoMos are also able to estimate possible mutation bias (Borges et al., 2022), which condition the genetic variability upon which selection can act. As such, PoMos can be helpful in testing the significance of biologically relevant processes responsible for molecular evolution, even in the absence of well-annotated reference genomes, as demonstrated by our application example on grasshopper species. Lastly, PoMos have been shown to be statistically sound; the maximum a posteriori tree is a consistent estimator of the species tree (Borges and Kosiol, 2020).

For all these reasons, PoMos constitute an attractive framework for estimating the species trees and testing the role of the different evolutionary forces responsible for molecular evolution across the genome. Here, we implement these models within the widely-used Bayesian programming tool RevBayes (Höhna et al., 2016) and demonstrate their application with population genomic data from a recent species radiation (Nolen et al., 2020).

### The model

PoMos model the evolution of a population of *N* haploid individuals and *K* alleles through time, in which changes in the allele content and frequency are both possible. These changes are mediated by evolutionary forces, such as genetic drift, mutation, and selection. The PoMo state-space includes fixed (or boundary) states {*N_a_i__*}. in which all *N* haploid individuals have the same allele *a_i_* ∈ {1,…, *K*}, and polymorphic states {*n_a_i__*, (*N* – *n*)*a_j_*}, in which two alleles *a_i_* and *a_j_* are present in the population with absolute frequencies *n* and *N* – *n*, respectively.

Mutations occur with rate *μ_a_i_a_j__* and govern the allele content, only arising in the fixed states:

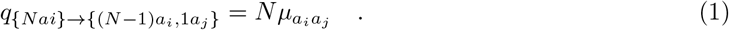

The assumption that mutations can only arise in the fixed states, corresponds to assuming that mutation rates are comparably low, and by the time of a new mutation a fixation of the previous mutation at the same site has already taken place. Low mutations rates are indeed verified for the majority of multicellular eukaryotes (Lynch et al., 2016). Additionally, a reversible mutation model is usually considered. In this case, we break the mutations into a base composition π and exchangeability parameter *ρ* (i.e., *μ_a_i_a_j__* = *ρ_a_i_a_j__π_a_j__*), similar to the definition of substitution rates in the general time-reversible model by Tavaré (1986). Assuming reversible mutations has no biological basis; however, this assumption simplifies obtaining formal quantities such as the stationary distribution while still allowing the modeling of mutation biases.

Genetic drift is modeled according to the Moran model, in which one individual is chosen to die, and one individual is chosen to reproduce at each time step. Selection acts to favor or disfavor alleles by differentiated fitness: *ϕ_a_i__* = 1 + *σ_a_i__*, where *σ_a_i__* is the selection coefficient of the *a_i_* allele. Together, genetic drift and selection govern the allele frequency changes:

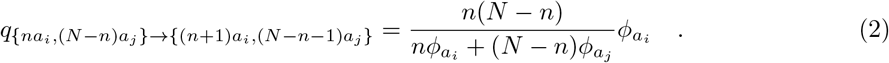

As standard substitution models, PoMos are continuous-time Markov models and are fully characterized by their rate matrices. PoMos express time in Moran generations, and the rates in equations (1) and (2) together define the PoMos rate matrices (Borges et al., 2022)

### PoMos implemented in RevBayes

Previously, PoMos were implemented in a Maximum Likelihood framework in HyPhy (De Maio et al., 2013) and IQ-Tree (Schrempf et al., 2019). Here, we implemented PoMos in a Bayesian framework in the phylogenetic software package RevBayes (Höhna et al., 2016) (**Fig. 1**). The main advantage of our new implementation are extensions of PoMos (described below) and the existing model toolbox of RevBayes, resulting in models whose parameters cannot be estimated in the Maximum Likelihood framework. RevBayes includes several maintained and extensively used routines for phylogeny estimation and hypothesis testing (Höhna et al., 2014, 2016, 2018, 2021). In addition, RevBayes can perform a range of inferences — e.g., phylogeny reconstruction, inference of selection, molecular dating, character evolution, ancestral state reconstruction — that can be easily married with PoMos, allowing the development of newly devised models for varied evolutionary applications.

**Figure 1:**
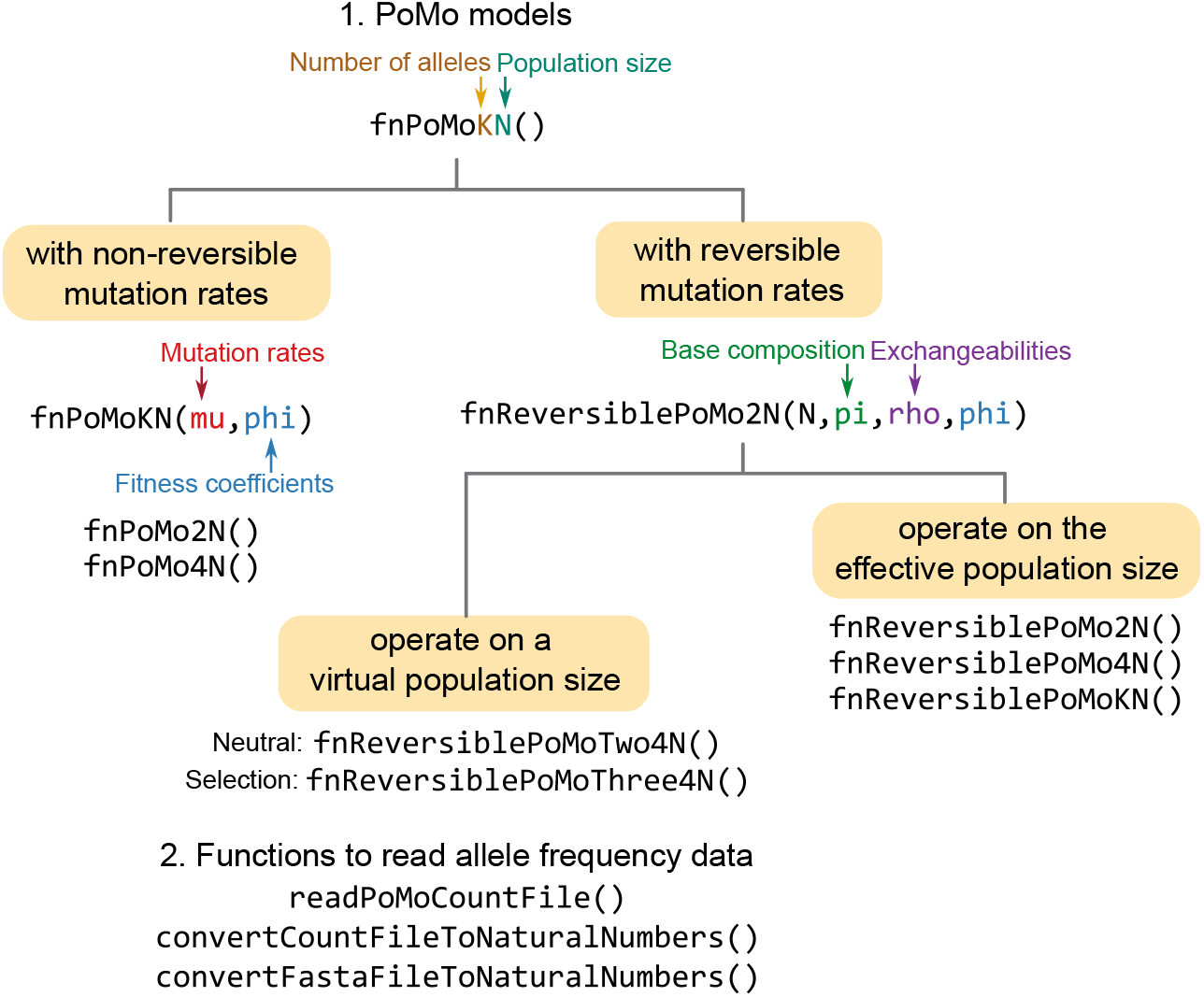
An overview of the PoMos functions implemented in RevBayes (Höhna et al., 2016).

PoMo rate matrices include selection by modeling allelic fitness (argument `phi`). Selection can also be helpful to model nucleotide usage biases that are not caused by selection but behave like it (Nagylaki, 1983). This is the case of GC-biased gene conversion, a mutational bias present in many living organisms (Galtier et al., 2009), that prefers GC alleles over AT during recombination. Neutral dynamics can simply be defined by assigning the fitness coefficients vector to **1**.

PoMo functions in RevBayes permit modeling mutational biases. Mutations can be assumed non-reversible (via the argument `mu`) or reversible (functions including the prefix `Reversible`). Reversible mutations assume that the mutation rate between any two alleles is the product of an exchangeability and a base composition term (arguments `rho` and `pi`). Another advantage of having PoMos in a Bayesian framework is that empirically measured mutation rates, population sizes, or nucleotide usage biases can be accounted for during inference via informative priors. Additionally, we implemented virtual PoMos, which were recently introduced (Borges et al., 2022): `fnReversiblePoMoTwo4N` and `fnReversiblePoMoThree4N`. Virtual PoMos mimic a population dynamic that unfolds in the effective population by using a much smaller virtual population size, making it computationally more efficient. While inferences are performed in the virtual population, the mutation rates and selection coefficients are rescaled to real dynamic using theoretically obtained scaling laws (Borges et al., 2022).

Apart from the standard PoMos, which usually operate with the four nucleotide bases, we have also added PoMo rate matrices to be used in genetic systems that might have any number of variants (i.e., `fnPoMoKN`). In addition, we included specific models for the biallelic case, as this is typically used in population genetic applications (e.g., `fnReversiblePoMo2N`).

Furthermore, we have included functions to read allele frequency data and correct for sampling biases. Genetic diversity is usually undersampled because sampled fixed sites might not necessarily be fixed in the original population. To account for this missing data, we implemented two corrections: the weighted and the sampled binomial corrections (described in Schrempf et al. (2016)). While the former directly uses the binomial weights to initialize the likelihoods of every PoMo state at the tree tips, the latter samples PoMo states from them. These methods are integrated into the functions that read allele frequency data, either from count files or FASTA alignments (i.e., `convertCountFileToNaturalNumbers or` `readPoMoCountFile`, and `convertFastaFileToNaturalNumbers` respectively).

### Implementation, availability and validation

The RevBayes source code is freely available at https://github.com/revbayes/revbayes. A detailed tutorial about Bayesian phylogenetic inference with PoMos is available at https://revbayes.github.io/tutorials/pomos/.

As a critical part of a robust Bayesian workflow, we validated our implementations of PoMos in RevBayes using a simulation-based calibration approach (Talts et al., 2018). This method assesses the soundness of a posterior sampler by testing the expectation that if data is generated according to a model, then the Bayesian posterior inferences with respect to that same model are calibrated by construction. In practical terms, if one draws *S* parameter values from their prior distributions and subsequently simulates S data sets using these parameter values, the inferred *α*% credible intervals will contain the true parameter value in *Sα* of times. This simulation-based calibration approach is quite stringent because the correct coverage of credible intervals is only achieved if the likelihood function, the MCMC sampler, and the simulation function are implemented correctly.

We simulated 1000 alignments of four species and 1000 sites, considering a population dynamic including three virtual individuals, four alleles (*A, C, G* and *T*) and general mutation (nine free parameters in total: three base composition parameters *π* and six exchangeabilities *ρ*) and selection (three fitness coefficients; *ϕ_A_* was set to 1) schemes. We calculated the coverage probabilities for each parameter and compared them with the credibility interval size. Our analyses show a 1:1 correlation (i.e., identity line) between the coverage and credible interval size, indicating that our implementations are statistically and computationally sound (**Fig. 2**). The simulated data sets and the RevBayes scripts and output files used in simulation-based calibration analyses are all provided as additional material (see Data availability statement for further details).

**Figure 2:**
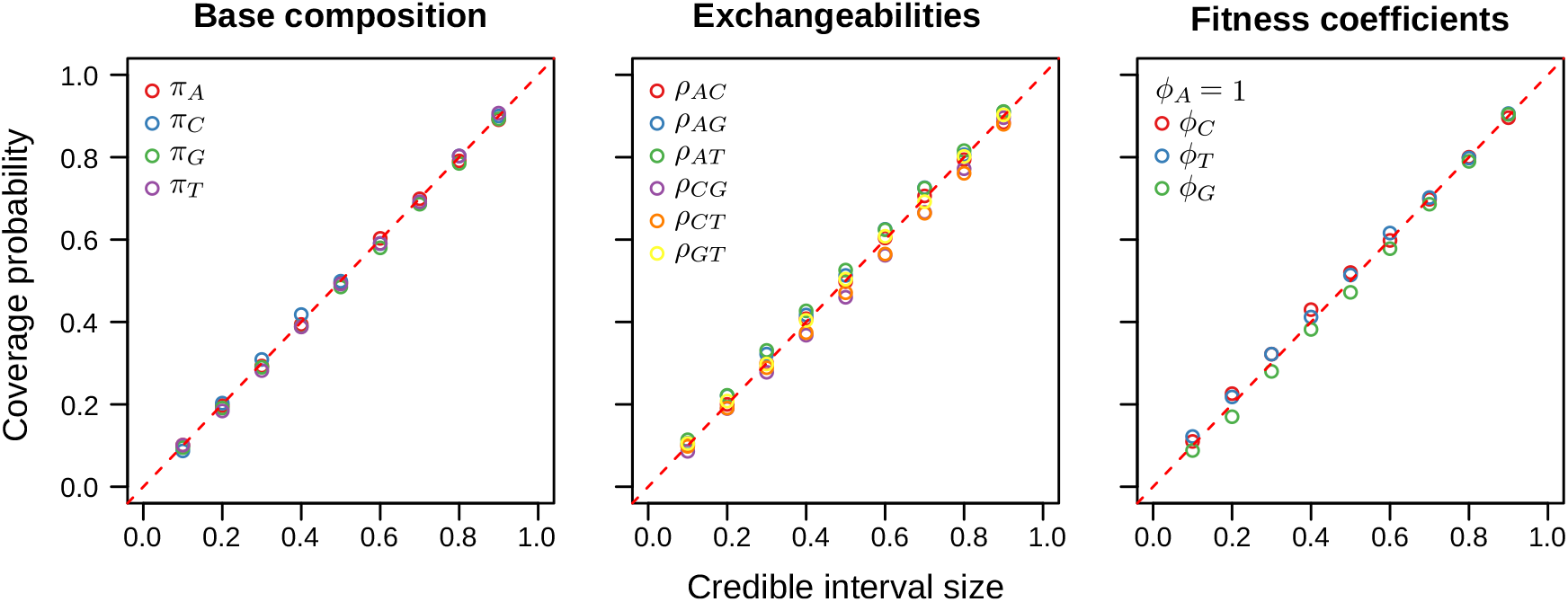
Simulation-based calibration of PoMos in RevBayes (Höhna et al., 2016) The plots were obtained using 1000 simulated data sets of multiple sequence alignments of 4 species and 1000 sites. We simulated a population dynamic including three virtual individuals, four alleles (*A, C, G* and *T*) and general mutation (nine free parameters in total: three *π*s and six *ρ*s) and selection (three fitness coefficients; *ϕ_A_* was set to 1) schemes. The coverage probability corresponds to the proportion of credible intervals that includes the true parameter value. The expectation from mathematical theory is that coverage and credible interval size show a 1:1 correlation.

### Application to grasshoppers transcriptome data

We studied the recent radiation of five grasshopper species using transcriptome data (Nolen et al., 2020) and PoMos in RevBayes. This data set includes nine grasshopper populations with five to 15 sampled individuals per population (**Fig. 3**). FASTA alignments from 1 895 protein-coding genes were generated by calling the most common base at each position so that each individual grasshopper is represented by one allele. As in the original study, the data used in the phylogenetic analyses included only ORFs from orthologous gene. Paralogous genes potentially involved in duplications were excluded from these analyses, as well as UTRs that can potentially be involved in miss-assemblies. Note that PoMos describe the evolution of haploid populations, so, as is usual in many population genetic models, the total number of individuals are the total of individual chromosomes (i.e., a population of N diploid individuals is seen as a population of 2*N* haploid individuals). Consequently, one should avoid making inferences with genomic units that are known to have a different number of individuals (e.g., heterosomes and sex chromosomes, or gene families with multiple paralogs, unless PoMos are properly combined with models of chromosome or gene family evolution).

**Figure 3:**
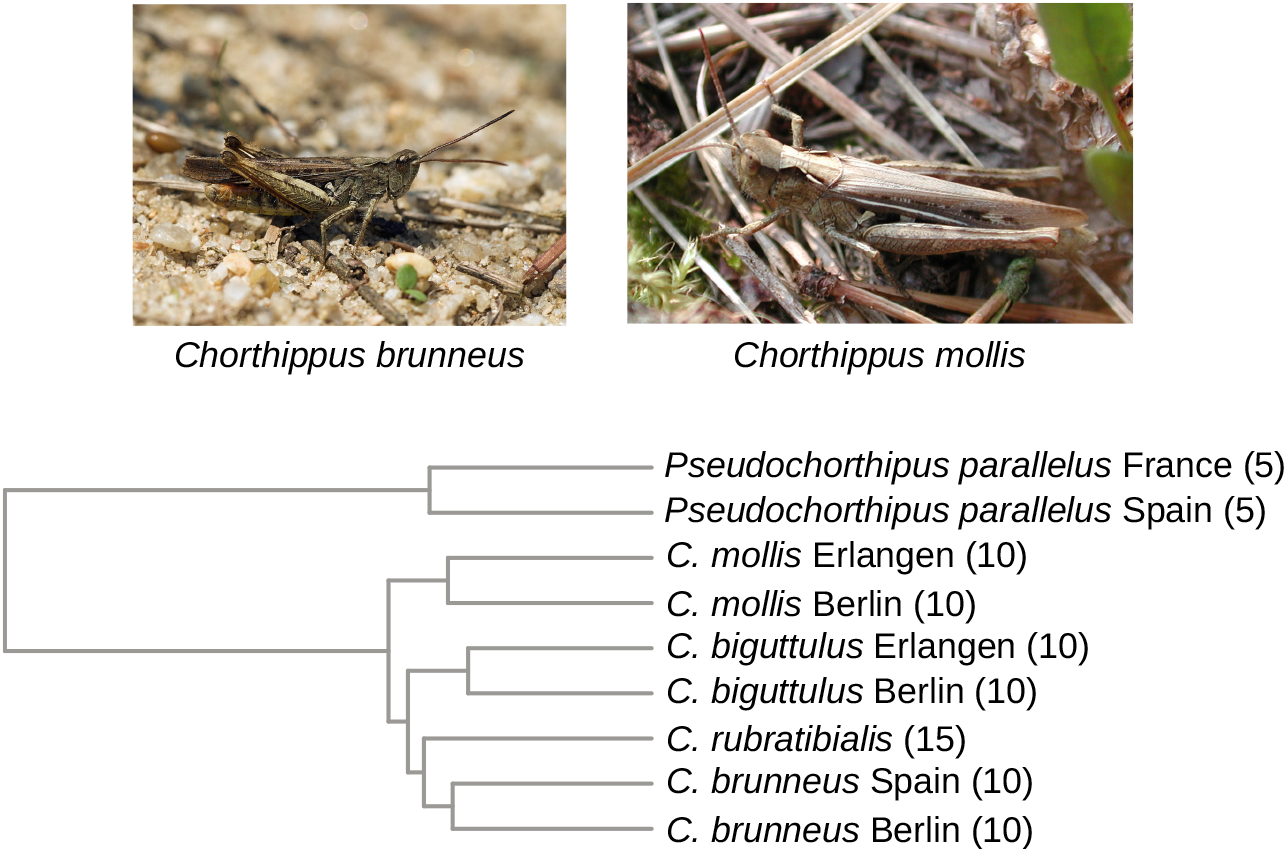
Grasshopper evolutionary history. Application example of the Bayesian PoMos in RevBayes to transcriptome data from nine grasshopper populations of the genus *Chorthippus sp.* (Nolen et al., 2020). Phylogenetic inferences were conducted using one million randomly sampled sites from the third codon position of 1 895 protein-coding genes together with the virtual PoMoThree model. The maximum a posteriori tree is depicted; all the branches had a posterior clade probability of 1.00 (not shown). The number of sampled individuals per population is indicated inside parenthesis. Photos of *Chorthippus brunneus* (credits to Jörg Hempel) and *Chorthippus mollis* (credits to G.-U. Tolkiehn) grasshoppers taken from Wikipedia under the Creative Commons Attribution-Share Alike 3.0 licence.

The final alignment included only the third codon position and those sites for which at least four individuals were observed per population. A total of 2 744 646 sites were obtained, but for computational reasons, only a sample of 1 million randomly selected sites was used for the phylogenetic inferences. This represents a manageable number of sites that are usually easily reached in standard empirical studies. We have repeated the phylogenetic analyses with two additional data sets of 1 million sites and obtained similar results that fully support the conclusions drawn here about the grasshoppers’ evolutionary history and pressures acting on their coding regions.

Phylogenetic inferences were carried out with the virtual PoMoThree model, which assumes a reversible mutation scheme. As we aimed to test the dynamic where AT alleles (or weak alleles, W) are preferred by mutational bias, and GC alleles (or strong alleles, S) are favored by GC-biased gene conversion, we defined the allele composition, exchangeabilities, and fitness coefficients in the following manner. The allele composition is generally defined as *π* = [*π_a_, π_C_, π_G_,π_T_*], which in terms of strong and weak alleles becomes *π* = [*π_W_, π_S_, π_S_, π_W_*]. The allele exchangeabilities are generally defined as *ρ* = [*ρ_AC_, ρ_AG_, ρ_AT_, ρ_CG_, ρ_CT_, ρ_GT_*], but we have only considered two exchangeability classes, one between weak and strong alleles, and another one between alleles of the same type: i.e., *ρ* = [*ρ_ws_, ρ_WS_, ρ_WW_, ρ_SS_, ρ_WS_, ρ_WS_*], where *ρ_WW_* = *ρ_SS_*. The fitness coefficients are generally defined as *ϕ* = [*ϕ_A_, ϕ_C_, ϕ_G_, ϕ_T_*], but because we are only interested in modelling GC-bias, we simplified it to *ϕ* = [1.0, *ϕ_S_, ϕ_S_*, 1.0], where *σ_S_* = *ϕ_S_* – 1 is the selection coefficient favouring the strong alleles.

In total, four parameters were estimated: *π_W_, ρ_WS_, ρ_SS_* = *ρ_WW_* and *ϕ_S_*. We set a Beta prior on *π*_W_ and an exponential prior on the exchangeabilities *ρ_WS_* and *ρ_SS_*. A reversible jump mixture was set on *ϕ_S_*, including a point mass at 1.0 and a gamma prior for all the positive values, both with probability 0.5. This means that the GC-bias rate can take on the constant value 0.0 (suggesting neutrality) or be drawn from the base distribution gamma (indicating selection). A special reversible-jump MCMC algorithm (for details see Freyman and Höhna, 2018) is then used to infer whether GC-bias was active or not. We assumed a strict molecular clock rate drawn from an exponential prior with an arbitrary root age that we fixed to 1.0. A uniform time tree prior was set for the phylogeny. We ran the MCMC with four chains for 300 000 generations. The Bayesian inferences in RevBayes with the grasshopper’s sequence alignment (one million sites and nine populations) took approximately 29 hours in an iMac desktop with a 3.7 GHz Intel Core i5 processor and 32GB memory using four parallel processes. Convergence was assessed via the effective sample size parameter, which was higher than 400 for the continuous parameters and tree splits (Fabreti and Höhna, 2021). The multiple sequence alignments, the RevBayes scripts and output files used in the grasshopper’s phylogenetic analyses are provided as additional material (see Data availability statement for further details).

The posterior distributions of the mutation rate parameters and selection coefficient, despite all showing clear peaks (**Fig. 4**), cannot be directly interpreted as they are relative to a virtual population size of three individuals. To obtain sensible estimates of these evolutionary forces, we need to scale them according to the grasshoppers’ effective population size. While empirical estimates of the genetic diversity (Watterson’s *θ* = 0.012, according to Nolen et al., 2020) are available for these species, we lack direct measurements of the mutation rate that would allow us to estimate their effective population size via *θ* = 4*Nμ*. We have instead used the fruit fly mutation rate (3.5×10^-9^, according to Keightley et al., 2014) and obtained an effective population size of approximately 850 000 individuals. We note that this estimate must be considered with care, as mutation rates vary greatly across phylogenetic scales.

**Figure 4:**
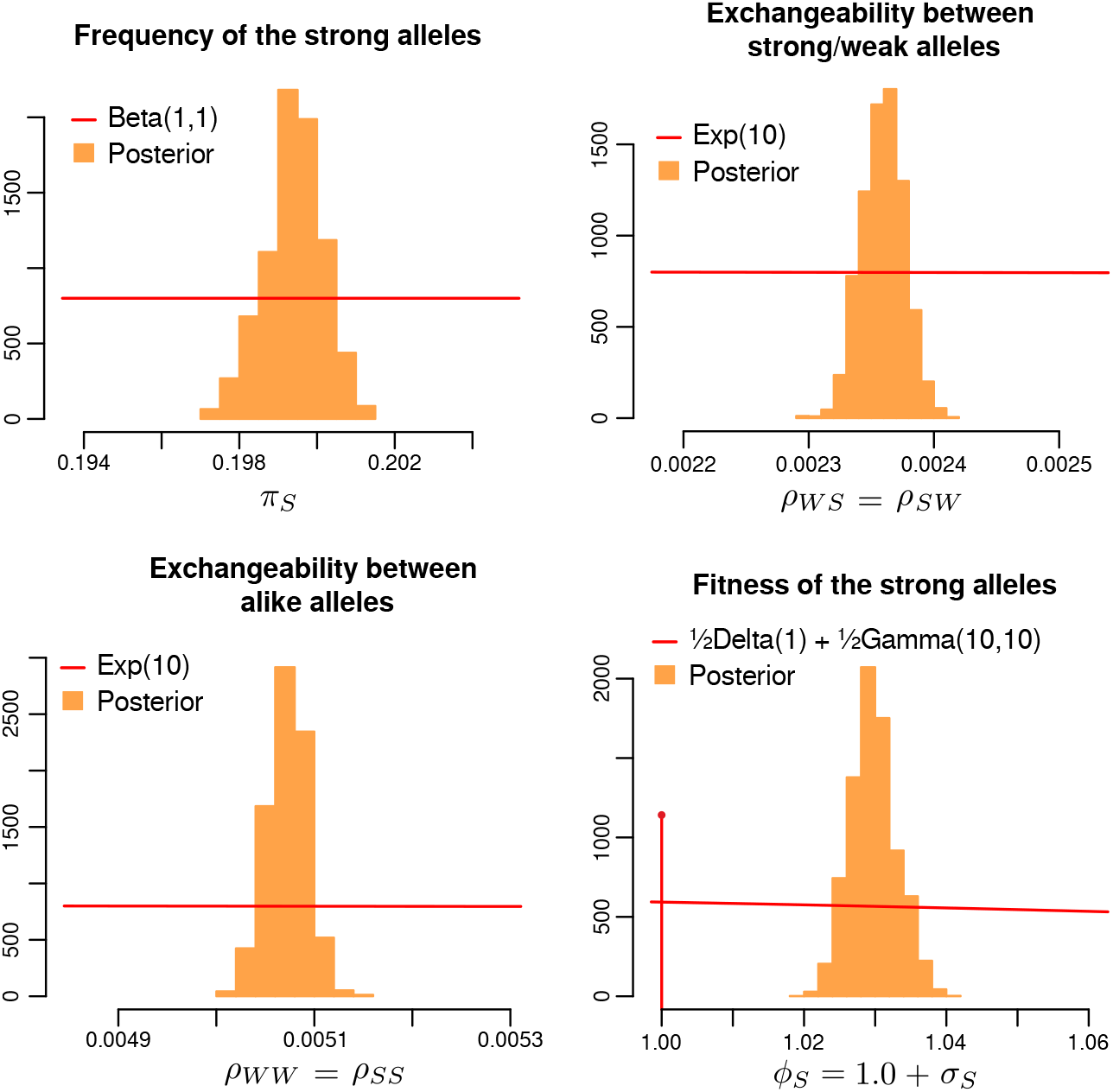
Posterior and prior distributions of the mutation biases and selection parameters at the grasshoppers’ third codon position. Delta indicates the mass point distribution. The prior densities are rescaled only to facilitate comparisons between the prior and posterior distributions and should not be read on the y-axis.

Using the estimated population size and the scaling laws in Borges et al. (2022), we calculated the relative mutation rates and GC-bias for the grasshoppers’ transcriptomic data (**Table 1**). Despite the four mutation types being around the same order of magnitude, we found evidence for mutational bias, with mutations from strong to weak alleles being more frequent than mutations from weak to strong alleles: *μ_SW_* = 5.67 × 10^-10^ and *μ_WS_* = 3.76 × 10^-10^. Mutations between alleles of the same type are the least frequent: *μ_WW_* = 2.64 × 10^-10^ and *μ_SS_* = 1.75 × 10^-10^. The GC-bias rate is two orders of magnitude higher than the mutation rates but two orders of magnitude lower than the reciprocal of the population size: *σ_S_* = 6.89 × 10^-8^ and *Nσ_S_* = 0.059. This indicates that GC-selection (or possibly GC-biased gene conversion) operates in a nearly neutral range. Nonetheless, the posterior of *ϕ_S_* never included the point mass 1.0 (log Bayes Factor = 3.68), indicating that selection in favor of GC alleles at the third codon position is very significant despite acting in the nearly neutral range.

**Table 1:**
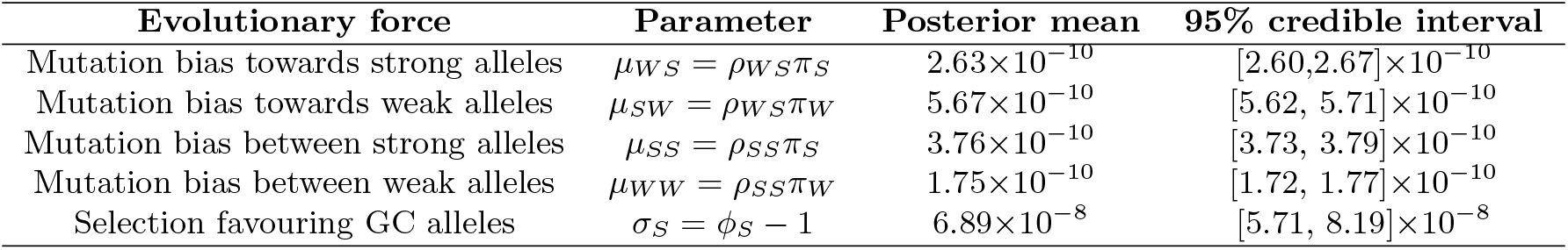
Posterior mean estimates and 95% credible intervals of the molecular dynamic parameters between weak (AT) and strong (GC) alleles at the grasshoppers’ third codon position. Note that 2*π_S_* + 2*π_W_* = 1, *ρ_SS_* = *ρ_WW_* and *ρ_SW_* = *ρ_WS_*. The estimates of the mutations rates and selection coefficients were rescaled for a population of 850 000 individuals based on the theoretical results of Borges et al. (2022).

Overall, these results suggest that a dynamic including mutational bias favoring AT alleles and selection bias promoting the fixation of GC alleles influences the codon usage patterns of the *Chorthippus* grasshoppers protein-coding sequences. Despite being reported here for the first time for the case of grasshoppers, these patterns have been observed in other taxa (e.g., Sueoka and Kawanishi, 2000), suggesting that gene GC-biased gene conversion is a general molecular process throughout the tree of life (Pessia et al., 2012; Lassalle et al., 2015).

Regarding the topology, our phylogenetic inferences show very well-supported branches [posterior probabilities (PP) all equal to 1.00 **Fig. 3**]. In agreement with Nolen et al. (2020), the populations of each species form very well supported clades. However, the relationships between *C. rubratibialis, C. brunneus* and *C. biguttulus* were unclear (PP = 0.67) in Nolen et al. (2020), whereas in our analyses the *C. rubratibialis* emerges as a sister species of *C. brunneus* (PP=1.0). Notably, the relative divergence among taxa is significantly larger than in Nolen et al. (2020). This result is expected in light of the mutation and selection bias we found acting on the grasshoppers’ coding sequences. We have recently shown that nucleotide usage bias leads phylogenetic methods to underestimate branch lengths (Borges et al., 2022). These biases are greater for more closely related populations as observed with these grasshopper populations. Indeed, *Pseudochorthippus* subspecies and *Chorthippus* species are estimated to have diversified recently, between 476 to 506 ka respectively (Nolen et al., 2020).

In this application, we show how PoMos can effectively help to characterize both the history and molecular evolution dynamic of non-model organisms with transcriptome data, which despite having well-known limitations compared to genomic data in the presence of well-annotated reference genomes, will continue to be used to understand the evolution of clades with large variation of genome size and structure (e.g. Irisarri et al. (2017); Hawlitschek et al. (2022)). However, PoMos have also been used with model organisms, such as the well annotated genomes of fruit flies (see Borges et al. (2022)). PoMos helped establishing possible routes of colonization of the African continent. By employing a phylogenetic molecular clock model, we dated the main episodes of fruit flies’ history such as their out of Africa migration. The advantage of having PoMos within RevBayes is that several means to solve the same problem exist. For example, one could have also dated the fruit flies phylogeny via informative priors on the mutation rates.

As it becomes easier and easier to sequence genomes and transcriptomes, we will soon have the resolution to solve long-standing questions in evolutionary biology that require leveraging intra-, interspecific, as well as phenotypic and ecological aspects of the evolutionary process. The combination of PoMos with phylogenetic models of molecular dating, species delimitation, phylogeography, and character evolution facilitate addressing this questions by scientist from broad backgrounds, in a user-friendly interface.

## Conclusion

We implemented PoMos into the widely-used phylogenetic Bayesian software RevBayes. Our implementation allows researchers to use the latest species tree inference models and combine state-of-the-art methods in phylogenetics to build robust inferences or for testing hypotheses. PoMos are able to perform accurate species tree inference while jointly estimating parameters of the molecular evolutionary dynamic. Together, PoMos offer more species tree inference methods that can be useful to characterize the evolution of both species and molecular sequences. We anticipate that our methods will be particularly interesting to researchers studying genomic data sets, including multiple species, populations, and individuals, which are increasingly being produced but still lack appropriate analytical methods. Furthermore, RevBayes is open-source and multiplatform; thus, it can easily be used on a variety of systems and settings for both academic research and teaching.

## Acknowledgements

This work was funded by the Vienna Science and Technology Fund (WWTF) [MA16-061] and partially supported by the Austrian Science Fund (FWF) [P34524-B] and Biotechnology and Biological Sciences Research Council (BBSRC) [BB/W000768/1]. This work was supported by the Deutsche Forschungsgemeinschaft (DFG) Emmy Noether-Program (Award HO 6201/1-1 to S.H.)

## Authors’ contributions

R.B. and C.K. conceived the project; R.B., B.B. and S.H. implemented the models in RevBayes; R.B. and R.P. performed the application example; R.B. and C.K. led the writing of the manuscript. All authors contributed critically to the drafts and gave final approval for publication.

## Data availability statement

The data used and produced by the the simulation-based calibration analyses of PoMos in RevBayes and the phylogenetic analyses conducted on the *Chorthippus sp.* grasshoppers (Nolen et al., 2020) are available at Zenodo (https://doi.org/10.5281/zenodo.6592395, last accessed May 30, 2022).

